# Automating literature screening and curation with applications to computational neuroscience

**DOI:** 10.1101/2023.12.15.571963

**Authors:** Ziqing Ji, Siyan Guo, Yujie Qiao, Robert A. McDougal

**Affiliations:** Biostatistics, Yale School of Public Health, Yale University, New Haven, CT, USA; Integrative Genomics, Princeton University, Princeton, NJ, USA; Biomedical Informatics and Data Science, Yale School of Medicine, Yale University, New Haven, CT, USA; Program in Computational Biology and Bioinformatics, Yale University, New Haven, CT, USA; Wu Tsai Institute, Yale University, New Haven, CT, USA

**Keywords:** Natural Language Processing, Large Language Model, Computational Neuroscience, Automated Curation

## Abstract

**Objective:** ModelDB (https://modeldb.science) is a discovery platform for computational neuroscience, containing over 1800 published model codes with standardized metadata. These codes were mainly supplied from unsolicited model author submissions, but this approach is inherently limited. We estimate we have captured only around one-third of NEURON models and lower fractions for other simulators. To more completely characterize the state of computational neuroscience modeling work, we aim to identify works containing results derived from computational neuroscience approaches and their standardized associated metadata (e.g. cell types, research topics).

**Materials and Methods:** Known computational neuroscience work from ModelDB and identified neuroscience work queried from PubMed were included in our study. After pre-screening with SPECTER2, GPT-3.5 and GPT-4 were used to identify likely computational neuroscience work and their relevant metadata.

**Results:** SPECTER2, GPT-4, and GPT-3.5 demonstrated varied but high abilities in identification of computational neuroscience work. GPT-4 achieved 96.9% accuracy and GPT-3.5 improved from 54.2% to 85.5% through instruction-tuning and Chain of Thought. GPT-4 also showed high potential in identifying relevant metadata annotations.

**Discussion:** Due to computational limitations, we only used each paper’s title and abstract, partially leading to false negatives. Further efforts should be devoted to including more training data and further improving current LLMs through fine-tuning approaches.

**Conclusion:** NLP and LLM techniques can be added to ModelDB to facilitate further model discovery, and will contribute to a more standardized and comprehensive framework for establishing domain-specific resources.

## INTRODUCTION

Over the years, numerous informatics resources have been developed to aggregate human knowledge, from generalist resources like Wikidata [1] to domain-specific scientific resources like GenBank [2] for nucleotide sequences and NeuroMorpho.Org [3] for neuron morphologies. Researchers are often obligated to submit nucleotide sequences to GenBank, but for many other types of scientific products, researchers have no such obligations, sometimes leading to sharing either not happening or not happening in a consistent manner (e.g. with standardized metadata). The broader scientific community, however, benefits most when scientific products are available in a consistent format.

Numerous domain-specific knowledge-bases attempt to address this need, but they face at least three major challenges: (1) identifying publications describing relevant research; (2) identifying relevant metadata for each scientific product; (3) obtaining additional needed details not present in the corresponding publication. Monitoring the literature is non-trivial; PubMed lists over 1.7 million publications in 2022. Even once the literature is filtered to a specific field, the scientific product and relevant metadata desired by knowledge-bases is often not explicitly mentioned in the title or abstract, so determining relevance of a given paper may require a careful reading of the full text. Traditionally, these challenges are addressed by human curators, but this requires both significant domain knowledge and time. Established repositories with community support may receive community contributions. On the one hand, individual researchers know their work best and are, for that reason, in the best position to identify relevant scientific products and metadata. On the other hand, contributing researchers are unlikely to be experts on the ontologies used by a specific repository. To streamline operations, some knowledge-bases have turned to using rule-based approaches (e.g. [4]) or Natural Language Processing techniques such as using a custom BERT-based model (e.g. [5]) to partly automate metadata curation, but these approaches often do not generalize well between knowledge-bases.

We describe a generalizable, cost-effective approach for identifying papers containing or using a given type of scientific product and assigning associated metadata. Our approach uses document-embeddings and citations of core pre-identified papers to systematically pre-screen publications. We use a large-language model (LLM) to confirm inclusion criteria and identify the presence-or-absence of categories of metadata in the abstract. For identified categories, the LLM is used to identify relevant metadata annotations from a list of ontology terms specified by the knowledge-base. We demonstrate the utility of this approach by applying it to identify papers and metadata for inclusion in ModelDB, a discovery tool for computational neuroscience research [6].

## MATERIALS AND METHODS

### Corpus acquisition

Our corpus includes 1564 abstracts (actually: titles and abstracts, but referred to as “abstracts” for convenience) for computational neuroscience models from ModelDB as well as generic neuroscience papers from 2022 extracted from PubMed with MeSH terms under either C10 (“Nervous System Diseases”) or A08 (“Nervous System”). Title and abstract of each paper were extracted. Our final corpus included a total of 105,502 neuroscience related abstracts from 2022.

### PCA general separability

To get a general sense of SPECTER2’s capabilities in identifying computational work, we implemented Principal Component Analysis (PCA) on a set of 3264 abstracts containing 1564 known computational models from ModelDB and 1700 generic neuroscience works, using their SPECTER2 embeddings [7]. This gives us a view of whether SPECTER2 embeddings contain information regarding whether a certain abstract is computational or not and how separable the two sets of abstracts are merely based on the embeddings.

### KNN-based approach with SPECTER2 embeddings

We then explored the vector space of SPECTER2 embeddings at a deeper level through looking at the k nearest neighbors within the known computational work from ModelDB. The SPECTER2 embeddings for both sets of abstracts are distributed within the same space for the purpose of exploring patterns of the distance to the kth nearest abstract (we considered k = 5, 10, 50, and 100). Specifically, we calculated two values: the fraction of *models* whose kth nearest model is within distance d in the set of models, and the fraction of *generic neuroscience works* whose kth nearest model is within distance d. The difference of these fractions would result in a maximum value and an associated distance, which is suggested to be an “optimal” distance value, for each k. Therefore, given an unknown abstract, if its kth nearest model is within the corresponding “optimal” distance, this abstract was deemed likely to include computational work.

With our test dataset, we recorded the distance to the kth nearest model for each abstract in the test dataset for different k values. The abstracts were then all sorted by this distance. Based on the result, we explored two hypotheses. Firstly, the ones with shorter distance values may have a higher possibility of being computational. Secondly, as k increases, more non-computational work may occur within the set of abstracts with smaller associated distances. To assess these hypotheses, for each k, the first 100 test abstracts with the smallest distances were annotated by two authors (SG, ZJ) to determine a gold standard for whether or not they include computational work.

### Prompt Engineering with GPT-3.5 and GPT-4

Works identified as likely to use computational neuroscience from the SPECTER2 embeddings were further screened by GPT-3.5 and GPT-4. Prompts were written take into consideration the distinction between computational neuroscience and models in other categories (ie. statistics models and biophysics models), as shown below:

*You are an expert in computational neuroscience, reviewing papers for possible inclusion in a repository of computational neuroscience. This database includes papers that use computational models written in any programming language for any tool, but they all must have a mechanistic component for getting insight into the function of individual neurons, networks of neurons, or of the nervous system in health or disease. Suppose that a paper has a title and abstract as indicated below. Respond with “yes” if the paper likely uses computational neuroscience approaches (e*.*g. simulation with a mechanistic model), and “no” otherwise. In particular, respond “yes” for a paper that uses both computational neuroscience and other approaches. Respond “no” for a paper that uses machine learning to make predictions about the nervous system but does not include a mechanistic model. Respond “no” for purely experimental papers. Provide no other output*.

*Title: {title}*

*Abstract: {abstract}*

GPT-4 was queried with the article title and abstract from each article in the acquired corpus with query specifying the task of classification. To analyze the implication from different input format and parameter values in classifying the extracted titles and abstracts to “Computational Neuroscience” and “Neuroscience”, performances between individual and concatenated query inputs were compared, while the generated outputs among temperature values of (t = 0, 1, 2) were analyzed. More efforts were later devoted to improving GPT-3.5’s results through Chain-of-Thought [8].

### Result evaluation through annotation agreement

The performances of SPECTER2, GPT-3.5, and GPT-4 were evaluated by comparing to ground truth, which is established through inter-annotator agreement (utilizing Cohen’s Kappa Coefficient). Due to the ambiguous separation between computational and non-computational work, two annotators with a background in health informatics (ZJ, SG) manually annotated 100 abstracts for each k value to determine ground truth. Annotations were performed in a rule-based manner, with initial agreement of conditions to screen computational or non-computational characteristics to eliminate human bias error. Protocols of annotations were established under the following conditions:

1. *The annotators go through the abstract and title of the individual paper to examine if any keywords or concepts related to computational neuroscience are present;*
2. *When computational keywords are present, the annotator take into considerations of the context of the paper to determine if such keywords and concepts are relevant, or if they are only mentioned for comparison purposes;*
3. *To reinforce the annotators’ decision, annotators also consider other sections of the paper as additional support, especially for the method section, where annotators are able to distinguish methods that may seem computational on the surface level but should not be considered, such as statistical models, metaanalysis, reviews and etc*.;
4. *After completing the entire process, annotators are asked to verify each other’s output results, in case of any misalignment of details that may affect the annotating performance;*
5. *Ground truth is established under the consideration of independent outputs from each annotator to evaluate the model performance*.

### Metadata identification

Apart from academic field classification for PubMed papers, we also assessed the positive predictive value of metadata identification as follows: GPT-4 was utilized to automatically identify metadata for 115 PubMed papers derived from k=5 group with the smallest distances, with queries containing terminology sources for paper concepts, regions of interest, ion channels, cell type, receptors, and transmitters contained derived from ModelDB’s terminologies. These prompts stressed that answers should only come from the supplied terminology lists, which GPT-4 mostly respected. (Attempts to use GPT-3.5 led to high numbers of off-list suggestions, and were not pursued further.) Additionally, GPT-3.5 was implemented first to perform such a task–however, with the large number of off-list suggestions as compared with the fixed terminology source from ModelDB, this approach was not pursued further.

In the meantime, to identify the accuracy of metadata identification from GPT-4, two annotators with a background in health informatics (AG) and biostatistics (YQ) performed manual validation to define discrepancy between paper presentations and GPT-4 identifications, neglecting the lack of precision of certain keywords resulting from the selection from ModelDB’s existing terminology.

Additionally, the annotation of metadata includes three categories to evaluate the model’s performance - “Correct”, “Incorrect”, and “Borderline”. While “Correct” and “Incorrect” explicitly determines the performance of the model, “Borderline” catches the keywords that are present in the paper, but are irrelevant to the main research of the paper, as well as the ones that are relevant to the concepts in ModelDB’s existing terminology, but not entirely correct given the granularity of concept. For instance, in “Dopamine depletion can be predicted by the aperiodic component of subthalamic local field potentials”, the cell type of “Dopaminergic substantia nigra neuron” is not explicitly mentioned in the abstract, but referenced to introduce into the focus of the paper [9].

## RESULTS

For this study, we restricted our attention to neuroscience papers published in 2022 as this is after the training cutoff date for GPT-3.5 and GPT-4 and has a defined end-date to allow completeness.

### Identification of candidate papers

As described previously, we included computational neuroscience models from ModelDB, and generic neuroscience papers from PubMed published in 2022. PubMed lists 1,771,881 papers published between January 1, 2022 and December 31, 2022. As 84.5% of models from ModelDB with MeSH terms contained entries in the C10 or A08 subtrees, we used the presence of these subtrees as a proxy for an article about neuroscience. We note, however, that 8% of the models in ModelDB were not associated with PubMed IDs. Using this criteria, we found 105,202 neuroscience-related papers in PubMed from 2022.

### PCA results

Applying PCA to the high-dimensional SPECTER2 embeddings of our two datasets -- 1,700 generic neuroscience abstracts and 1,564 computational neuroscience abstracts -- revealed a notable separability in SPECTER2 embeddings between the two datasets (Figure 1). Thus SPECTER2 embeddings can capture relevant information pertaining to computational aspects of scientific abstracts within the broader neuroscience corpus.

**Figure 1.**
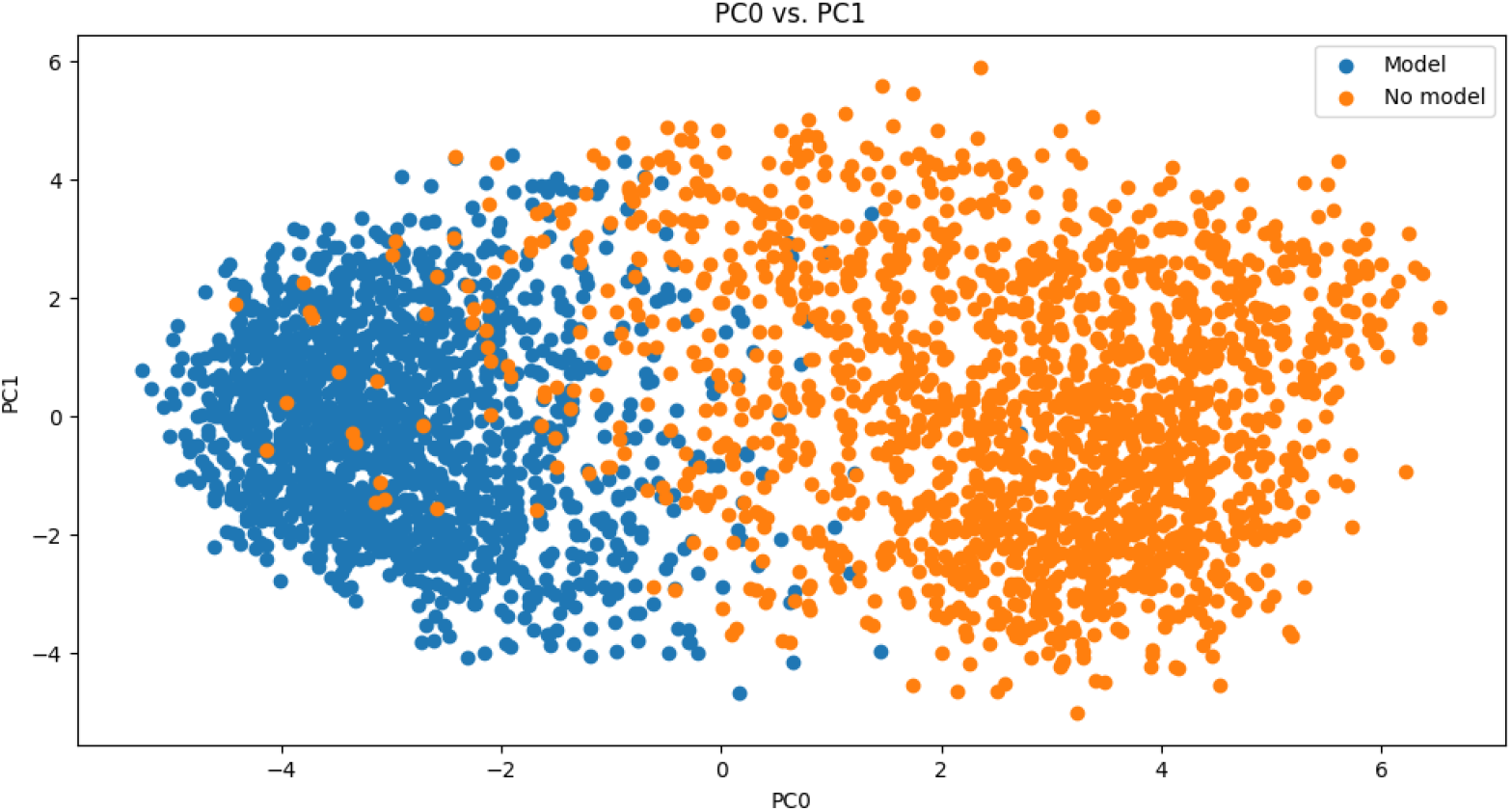
PCA analysis of the SPECTER2 embeddings exhibits clear separation between modeling (blue) and non-modeling (orange) along the 0th principal component.

### SPECTER2 vector space exploration

Based on the SPECTER2 embeddings of 1,564 known models from ModelDB and 1,700 generic neuroscience models, we implemented a knn-based approach in the vector space containing all the embeddings to examine patterns in their distribution and to determine how likely a neuroscience abstract is computational or not.

Given our hypothesis that computational modeling papers embed near other computational modeling work, we examined the differences between percentages of models (Figure 2A) and non-models (Figure 2B) with kth nearest known model within a certain distance. We used this difference (Figure 2C) to determine a cutoff threshold for when a paper should be considered likely to involve computational neuroscience work (i.e., if the kth nearest model is within the threshold). From this, we generated a possible range in which this threshold distance lies. For example, when we consider 5 nearest neighbors (k = 5), it is highly likely that a neuroscience work is computational when its 5th nearest neighbor (among the set of known models) is within the distance of approximately 0.08. For different k values, the peak distance is roughly different. As k increases, the value where the peak distance occurs increases as well. Different values of k influences the accuracy of results. As the value of k increases, more neighbors are taken into consideration, which can create bias in results and produce false positives or false negatives.

**Figure 2.**
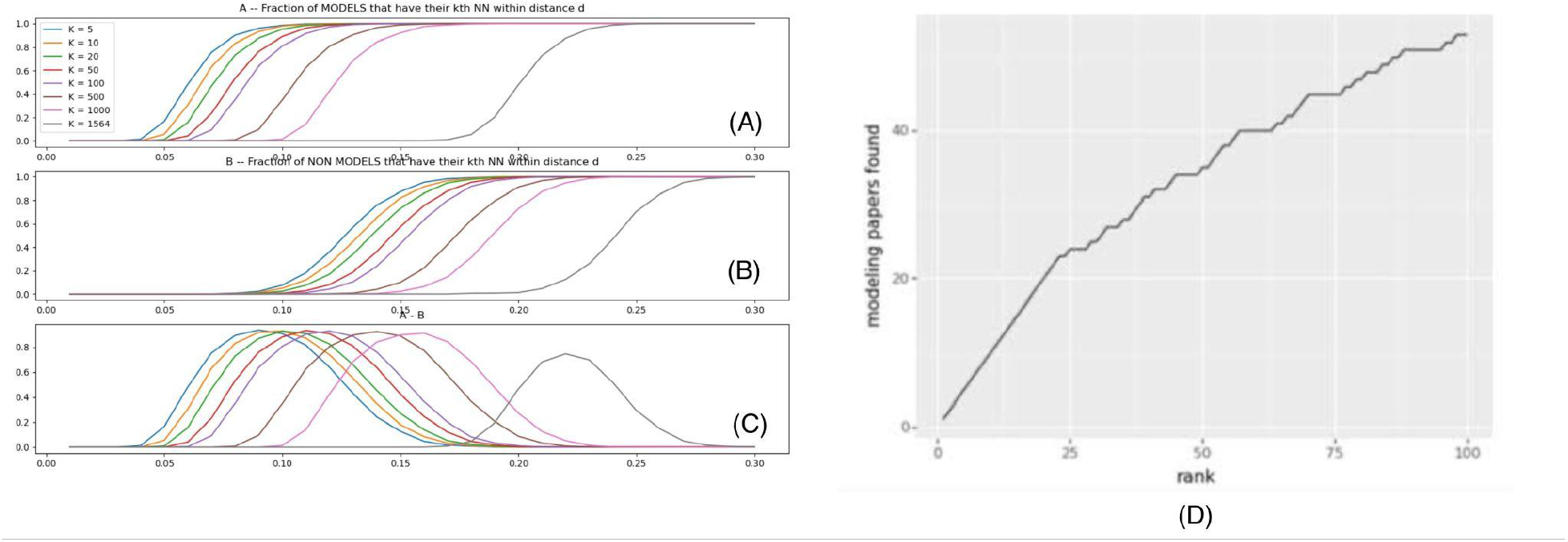
SPECTER2 Analysis. (A) The calculated percentages of known models that have their kth nearest neighbor (a known model) within a certain distance. (B) The percentages of generic neuroscience abstracts that have their kth nearest model within a certain distance. (C) The difference between A and B. (D) Computational Work from SPECTER2 Results (k=5)

With these potential thresholds, we explored the vector space of our test dataset, containing 105,502 neuroscience-related abstracts published in 2022. For different values of k, highly potential computational abstracts were identified and sorted by the distance to their kth computational neighbor. Two annotators manually annotated the top 100 abstracts for k = 5, 20, and 50 to determine the accuracy levels. Results show that k with smaller values generate more accurate predictions of computational characteristics of an abstract. Taking k = 5 as an example, as shown in Figure 3, the top 20 abstracts identified by SPECTER2 are all computational models. Going further down the list, more non-computational works start to emerge.

**Figure 3.**
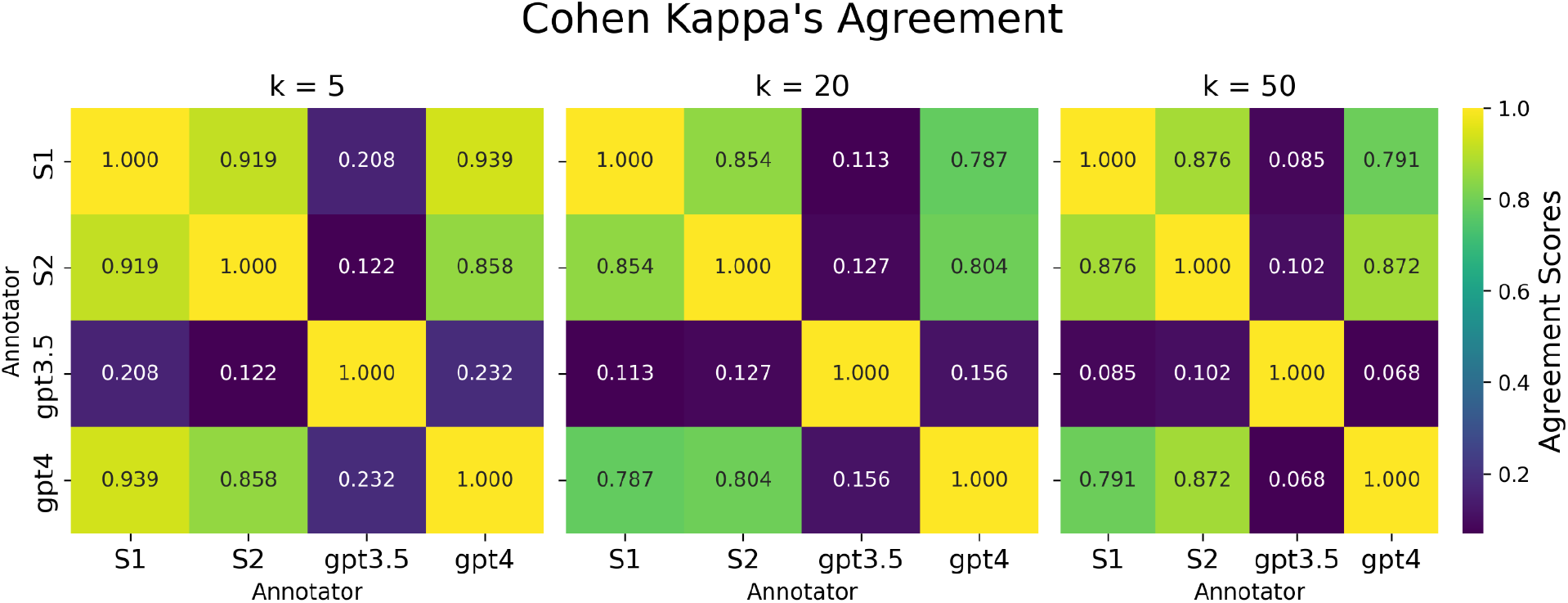
Cohen Kappa’s Agreement

### GPT-3.5 and GPT-4 results

We queried both GPT-3.5 and GPT-4 to find out if they would identify each paper -- given the title and abstract -- as using computational neuroscience work or not. This check was performed using the prompt described in methods but without examples for the top 100 abstracts from SPECTER2 analysis results with k = 5, 20, and 50. These prompts were initially run with temperature = 0 for near-deterministic results. Cohen Kappa’s agreement index between the two annotators, as well as agreement with GPT-3.5 and GPT-4 is shown in Figure 3. GPT-3.5 tends to have much lower agreement scores with human annotators, compared to GPT-4. We also notice that GPT-3.5 often misclassified non-computational work as computational, leading to false positives.

### Improving GPT-3.5 Results

We employed three strategies to improve the F-1 performance of the relatively low-cost GPT-3.5 model:

1. Instruction Fine-tuning: Allowing Outputs of Uncertainty

As GPT-3.5 results returned a large number of false positives, we tweaked the prompt to allow GPT-3.5 to output unsure when it is not certain about the computational characteristics of an abstract. All instances classified as “unsure” are actually non-computational, according to GPT-4’s outputs. Integrating “unsure” instances into the negative set increased GPT-3.5’s F-1 score to 61.7%.

1. Role of Temperature

Using the fine-tuned prompt from the previous step, we further explored the role of temperature by using temperature (randomness) values 0, 0.5, 1, 0.5, and 2. The best performing temperatures as measured by F-1 scores were 0.5 and 1, as measured by a majority vote of three calls to the API (Figure 4).

**Figure 4.**
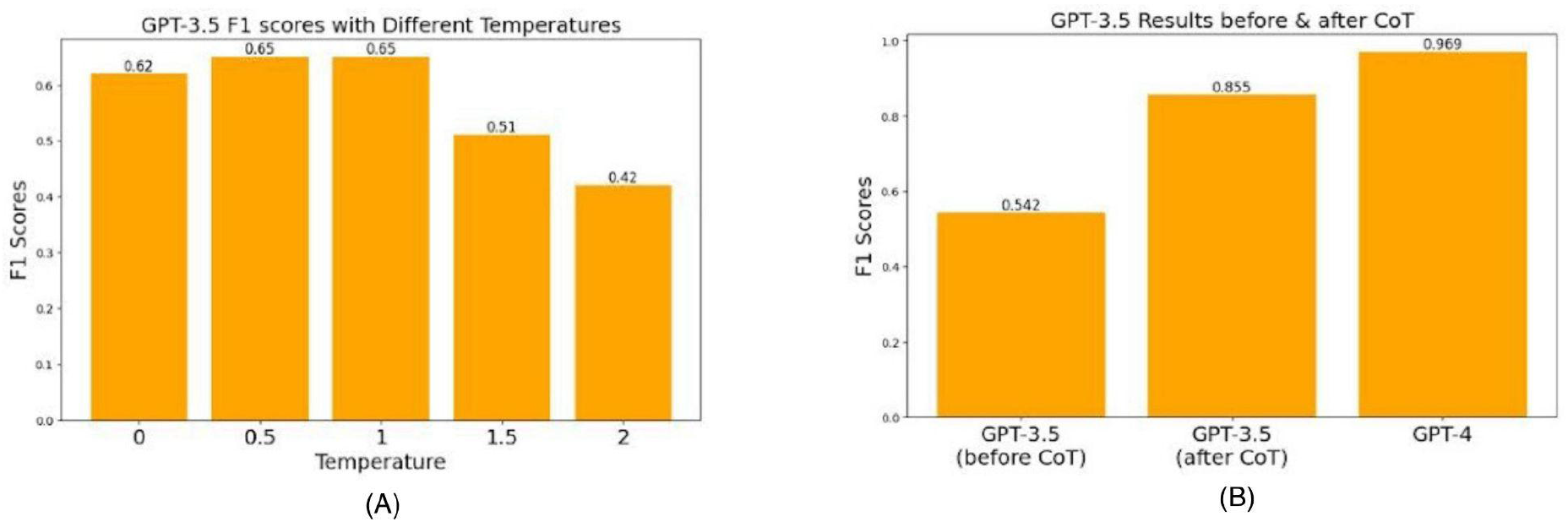
(A) Role of Temperature. GPT-3.5 was queried 3 times for whether or not an abstract implied use of a computational model. After combining the answers by voting, moderate but low temperature (randomness) achieved the highest F-1 scores, with high temperatures showing poor performance. (B) GPT-3.5 F-1 scores before and after CoT, compared with GPT-4

1. Chain-of-Thought Prompting

Following [8], which showed that incorporating reasoning steps (known as Chain-of-Thought) in LLM prompts can improve their performance, we added reasoning steps of how a human would approach classification tasks of computational work. GPT-3.5’s F-1 score, even though still lower than GPT-4, showed a marked increase to 85.5% after Chain-of-Thought was applied (Figure 4).

### Metadata Identification

We used repeated queries to GPT-4 to identify relevant computational neuroscience related metadata annotations for each remaining abstract. Specifically, we performed separate queries for each abstract for ModelDB’s long-standing six broad categories of metadata: “brain regions/organisms”, “cell types”, “ion channels”, “receptors”, “transmitters”, and “model concepts”. Each category may have hundreds of possible values and models may have multiple relevant metadata tags for a given category; for example, a model that examines the concept “Aging/Alzheimer’s” may also study the concept “synaptic plasticity.” 78.26% of the returned metadata annotations from applying GPT-4 to 115 abstracts were correct, as assessed by manual review. In exploratory studies, GPT-3.5 was relatively prone to suggest new terms or reword terms and so we did not apply it here.

Figure 5 illustrates the distribution of metadata tags assigned by the GPT-4 query to 115 abstracts, totaling 1,154 tags across various categories. Notably, the category “model concepts” received the highest number of tags, amounting to 753. It’s important to note that since paper titles and abstracts do not encompass all details about a model, our evaluation focused solely on the relevance of the predicted metadata tags, rather than the comprehensiveness of their model descriptions.

**Figure 5.**
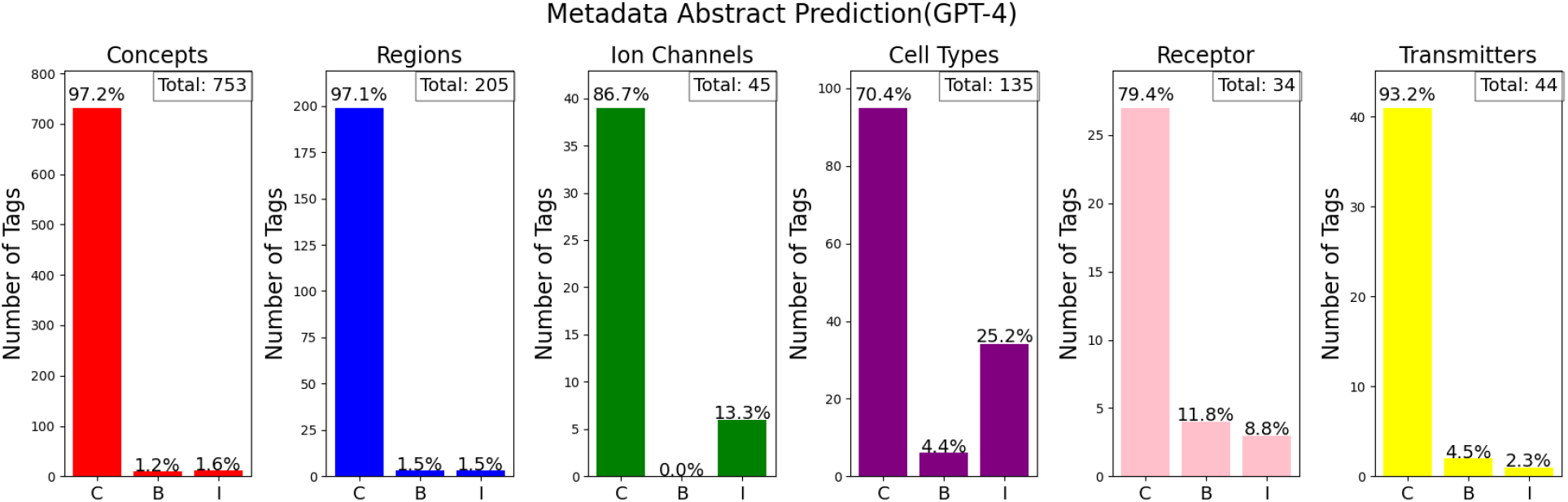
Metadata Identification Accuracy Across Categories

To ensure the accuracy of metadata, we relied on the consensus of two annotators for validation. The assessment across six categories revealed that “‘model concepts” achieved the highest prediction accuracy at 97.2%, while “cell types” recorded the lowest at 70.4%. These findings indicate that our approach, which leverages GPT-4 for metadata identification, shows comparable effectiveness to the rules-based approach [4], with the added benefit of circumventing the need for developing an explicit set of rules.

## DISCUSSION

Despite ongoing efforts (e.g. NIFSTD; [10]), there remains a lack of widely used robust and consistent ontologies in neuroscience ([11,12]). MeSH terms used to index publications in

PubMed are focused on medically relevant terms, missing the cellular-level details essential for characterizing neuroscience. Therefore, establishing an automated process for identifying and annotating computational neuroscience work and building efficient knowledge repositories will enhance reimplementation of computational models and exploit more potential in research scopes. In the future, integrating NLP techniques and LLMs may play an essential role in determining a publication’s computational aspects and metadata extraction.

### Using only ModelDB data and why knn-based approach?

We initially used entries from ModelDB [6] to generate SPECTER2 embeddings to establish an initial vector space of computational neuroscience work. However, ModelDB is not necessarily representative of computational neuroscience as a whole [13]. An alternative approach for creating this initial set is to collect abstracts from journals that are deemed as most relevant, such as Journal of computational neuroscience etc. However, only a small fraction of ModelDB models are in these journals and many computational models are paired with experimental studies. With this initial set of known computational work, we designed a KNN-based approach mainly since it is hard to generate a corresponding negative set that consists of generic neuroscience work with completely no computational work. Additionally, taking into account the nonlinearity of the vector space, alternative approaches like SVM, considering SVM’s struggle to define optimal linear boundaries, will not generate ideal results.

### SPECTER2 embeddings are only from title and abstract

We used SPECTER2 [7] document-level embeddings to determine whether a publication counts as publication work. A unique feature of SPECTER2 is that it was trained on citations but does not need them to produce an embedding. In our case, if the publications that cite or are cited by a publication are computational, it is more likely that this publication is also computational. Citation links provide more information on the publication’s content and explicitly using them would generate embeddings with higher representativeness. Another challenge is that the abstract does not necessarily focus on key model details (e.g. model 87284 [14] uses CA1 pyramidal cells, which are not mentioned in the abstract). This is due to several reasons, including permission to access full text, limited memory for processing and generating embeddings, and potentially overfitting by including too much information at the same time. Covering too much information in embeddings could lead to false positives in the results [4].

### What counts as computational neuroscience work?

Challenges encountered during manual annotation mainly revolved around deciding what exactly counts as computational work. Computational neuroscience [15] is not entirely well defined. There exists different opinions on what kinds of model or on what scale does the publication land within the scope of computational neuroscience. As an interdisciplinary field, there are a lot of methodologies when approaching a research question, which also indicates that it is hard to establish an explicit boundary that distinguishes between computational and non-computational work in the neuroscience discipline. Therefore, we defined a scope when prompting responses from GPT-4 and GPT-3.5, and the two annotators also followed the same standard as stated in the prompt during the annotation process.

### Metadata

When evaluating metadata predictions, annotators mainly considered the relevance of the provided tags to the papers themselves, while acknowledging the circumstances where certain tags could be mentioned, but to only serve as a reference or contrast. Borderline evaluation of the models’ results are mainly resulted from misalignment of terminologies from the ModelDB existing terminologies and the actual context mentioned in each paper. For example, when papers mention “globus pallidus” as the primary neurotransmitter, and GPT-4 outputs “Globus pallidus principal GABA cell” as the identified cell type given that such term has the closest meaning as compared to the actual context, annotators would evaluate such situation as “Borderline”.

## LIMITATIONS

### Using one-shot/few-shot learning

As shown in the analysis of GPT-3.5 results, subtle changes in prompts may yield large differences in the results. In this study, prompts given to GPT-3.5 and GPT-4 did not include concrete examples and expected results. Providing examples might improve results. Research on Few-Shot Learning (FSL) shows that models can learn to generalize information from only a limited number of examples and perform with high accuracy on a new task [16]. For example, in our study, providing a few examples of abstracts with their associated metadata might result in better results from GPT-3.5 and GPT-4. Future endeavors could potentially devote to experimenting with One-Shot Learning (OSL) or FSL, and measure the improvement in results.

### Fine-tuning LLMs for Metadata Extraction

Achieving state-of-the-art results in extracting metadata information in computational neuroscience might require further fine-tuning a LLM. Fine-tuning a LLM on a specific task can markedly improve performance [17], but to employ this in the curation context will require the development of a large high-quality dataset of examples.

## CONCLUSION

Recognizing the increasing importance of repository curation and how this could benefit future relevant work in neuroscience, we explored the feasibility of leveraging Natural Language Processing (NLP) techniques and Large Language Models (LLMs) to enhance identification and metadata extraction of computational neuroscience work.

SPECTER2 embeddings yield promising outcomes, demonstrating high-effectiveness for screening large numbers of neuroscience papers for the use of computational models without incurring API charges. GPT-4 shows relatively high accuracy in identifying metadata, but requires continued exploration in metrics to increase performance. Automating the process of collecting computational work and systematically storing information related to computational models paves the way for future research to reuse, update, or improve existing models.

Continued efforts should focus on increasing the accuracy of LLMs in gaining more accurate and contextually aware results. Attempts at using alternative metrics to improve accuracy of LLMs or establishing a new dataset comprising manually labeled computational information and metadata to fine-tune current large language models can potentially help reach state-of-the-art results.

Overall, this study marks a significant step forward in exploring the potential of NLP techniques and LLMs in automating the identification and curation of computational neuroscience work.

